# ColocZStats: A Z-Stack Signal Colocalization Extension Tool for 3D Slicer

**DOI:** 10.1101/2024.06.23.599080

**Authors:** Xiang Chen, Teena Thakur, Anand D. Jeyasekharan, Touati Benoukraf, Oscar Meruvia-Pastor

## Abstract

Confocal microscopy has evolved as a widely adopted imaging technique in molecular biology and is frequently utilized to achieve accurate subcellular localization of proteins. Applying colocalization analysis on image z-stacks obtained from confocal fluorescence microscopes is a dependable method to reveal the association between different molecules. In addition, despite the established advantages and growing adoption of 3D visualization software in various microscopy research domains, there has been a scarcity of systems supporting colocalization analysis within a user-specified region of interest (ROI). In this context, several broadly employed biological image visualization platforms were meticulously explored in this study to comprehend the current landscape. It has been observed that while these applications can generate three-dimensional (3D) reconstructions for the z-stacks and in some cases transfer them into an immersive Virtual Reality (VR) scene, there is still a lack of support for performing quantitative colocalization analysis on such images based on a user-defined ROI and thresholding levels. To address these issues, an extension called ColocZStats has been developed for 3D Slicer, a widely used free and open-source software package for image analysis and scientific visualization. With a custom-designed user-friendly interface, ColocZStats allows investigators to conduct intensity thresholding and ROI selection on imported 3D image stacks. It can deliver several essential colocalization metrics for structures of interest and produce reports in the form of diagrams and spreadsheets.

## 1 INTRODUCTION

Compared to conventional fluorescence microscopes, the most significant advantage of confocal microscopes is that they can exclude the out-of-focus light from either above or below the current focal plane (Jonkman et al., 2020). This capability facilitates precisely detecting the specific organelle in which the target molecule is present. In addition, the confocal microscope’s features of sharpening fluorescence images and reducing haze contribute to enhancing image clarity (Collazo et al., 2005). Many of these two-dimensional (2D) image slices can be continuously collected from focal planes at different depths along the z-dimension and eventually assembled to produce a z-stack comprising the specimen’s entire 3D data (Collazo et al., 2005; Theart et al., 2017).

Examining interactions between different proteins or molecular structures holds significant importance in biological sciences. Biologists often perform colocalization analysis on confocal image z-stacks to better understand the roles and interactions of proteins (Pompey et al., 2013; Bolte and Cordelières, 2006). Colocalization detects the spatial overlap between distinct fluorescent labels with different emission wavelengths to determine whether the fluorophores are close or within the same region (Lacoste et al., 2000; Adler and Parmryd, 2013). Colocalization involves two aspects: co-occurrence and correlation. Co-occurrence refers to the simple spatial intersection of different fluorophores. Correlation refers to distinct fluorophores codistributed proportionally with a more apparent statistical relationship (Adler and Parmryd, 2013; Dunn et al., 2011). Typical application examples of colocalization analysis include confirming whether a specific protein associates with microtubules (Bassell et al., 1998; Nicolas et al., 2008) or mitochondria (Lynch et al., 1996), or verifying whether different proteins associate with identical plasma membrane domains (Lachmanovich et al., 2003).

Visualizing superimposed fluorescence micrographs is the most common method for assessing colocalization. When the images of each fluorescence label are merged, their combined contribution can be indicated by the color of the microstructure appearance. For instance, because of the combined effects of green and red fluorescence, the colocalization of fluorescein and rhodamine can be recognized in yellow structures (Dunn et al., 2011). When analyzing z-stacks, a popular visualization method involves performing 3D reconstruction using techniques such as volume rendering, transforming the data into 3D semi-translucent voxels to enhance the user’s perception when observing samples (Theart et al., 2017; Lucas et al., 1996; Liu and Chiang, 2003). Another cutting-edge visualization technology is Virtual Reality (VR). VR is a digitally created immersive 3D simulated environment that closely resembles reality. While being fully immersed in this environment, users can navigate through and interact with virtual objects. It is noteworthy that as VR technology has developed, the characteristic it supplies that allows immersive observation of biological samples has led to this technology being continuously combined with a variety of visual analysis methods in recent years, enabling biologists to obtain a more realistic 3D awareness in the process of scientific exploration (Patil and Batra, 2019; Sommer et al., 2018). Although the application of VR in various subfields of bioinformatics has been steadily increasing, a study published in 2017 by Theart et al. (Theart et al., 2017) revealed that, before that time, no applications had provided the capability for colocalization analysis in VR. Nevertheless, this study, along with its subsequent research (Theart et al., 2018), collectively demonstrated the substantial potential and advantages of performing colocalization analysis for z-stacks based on the generated volume-rendered images in an immersive VR environment, namely that the efficiency of conducting such analysis and the precision of inspecting and assessing biological samples can be significantly improved.

In consideration of the above background, several well-known applications, with a focus on 3D graphics systems with VR functionalities, were investigated during the study period to gain an overview of the current status of these commonly accepted bioimaging visualization and analysis platforms for visualizing confocal microscopy data and evaluating its degree of colocalization.

ExMicroVR (Immersive Science LLC., 2024b) is a VR tool created for the immersive visualization and manipulation of multi-channel confocal image stacks. Its VR environment and easy-to-use user interface allow it to considerably expand microscopic samples so biologists can view and explore the molecules’ structures in greater detail. Another software, ConfocalVR (Immersive Science LLC., 2024a), in its current form, is an upgraded version of ExMicroVR. Not only does it have more added interactive features, but it also provides a range of relatively advanced image analysis capabilities, such as adding markers in 3D scenes, counting the number of interesting objects, and measuring the distance between them, allowing researchers to further investigate the complexity of cellular structures. ChimeraX (Goddard et al., 2018; Pettersen et al., 2021) is an interactive platform for visualizing diverse types of data, including atomic structures, sequences, and 3D multi-channel microscopy data. It supplies around 100 different analysis functionalities. ChimeraX VR is the VR extension of ChimeraX, enabling users to interact with cellular protein structures with stereo depth perception. When activated, a floating panel, precisely the same as the desktop interface, is available in the VR scene to help users use controllers to perform all necessary manipulations on the images. 3D Slicer (Fedorov et al., 2024) is a broadly recognized, accessible, and open-source platform that provides multifarious biomedical image processing and visualization features (Kikinis et al., 2013). Similarly, SlicerVR is the VR extension within 3D Slicer. Thanks to the capabilities of the 3D Slicer ecosystem, SlicerVR offers seamless VR integration in this popular image computing application (Lasso et al., 2018) so that observers can quickly transfer images displayed in desktop mode to the VR scenario. Additionally, it supports multi-user collaboration, allowing images within the exact scene to be manipulated synchronously (Pinter et al., 2020).

Through this investigation, it was found that all of the platforms above are capable of reading datasets generated by fluorescence confocal microscopy and converting them into 3D semi-translucent voxels that can be delivered to VR scenes for visualization. For instance, ConfocalVR offers a ‘blend function’ option that allows the extraction of voxels representing the spatially intersecting regions between two channels. The 3D rendering of these voxels can aid researchers in visually assessing the level of colocalization between channels. However, for more comprehensive colocalization analysis, relying solely on subjective identification of the relative distribution of different molecules from a visual perspective is insufficient. Objective and quantitative analysis is crucial. Despite the commendable image visualization and diverse analytical capabilities inherent in these platforms, there is still room for them to improve in obtaining colocalization statistics for confocal stacks. As indicated by a published work of Stefani et al. (Stefani et al., 2018), developing a dedicated tool to gain objective colocalization measurements remains one of the goals and challenges for ConfocalVR.

To address the aforementioned limitation and take advantage of the comparatively superior high extensibility of 3D Slicer, which supports the creation of interactive and batch-processing tools for various purposes (Kapur et al., 2016), a free, open-source extension called ColocZStats has been developed for 3D Slicer. ColocZStats is currently designed as a desktop application, which enables users to visually observe the spatial relationship between different biological microstructures while performing thresholding and ROI selection on the channels’ 3D volumetric representations via an easy-to-use graphical user interface (GUI) and then acquiring critical colocalization metrics with one-mouse click. The proposal of this tool contributes to supplementing the capabilities of 3D Slicer in visualizing multi-channel confocal image stacks and quantifying colocalization, thereby broadening its scope as an integrated image analysis platform.

## 2 MATERIALS AND EQUIPMENT

### 2.1 Description of Sample Data Source

To showcase the capabilities of ColocZStats discussed in this paper, we utilized confocal z-stack data collected during a study on the colocalization of DSS1 nuclear bodies with other nuclear body types. Please refer to the document/Supplementary Materials for a detailed description of the image. DSS1, also known as SEM1, is a gene that encodes a protein crucial for various cellular processes, most notably the function of the 26S proteasome complex in protein degradation. More specifically, a human ovarian clear cell carcinoma cell line (RMG-I) was seeded at 100,000 cells per well onto a coverslip in a 6-well plate and allowed to grow till 70% confluency. On the day of staining, the cells were fixed with 4% ice-cold paraformaldehyde for 15 mins at room temperature (RT) and blocked with 2% Bovine Serum Albumin (BSA) in 0.1% Phosphate Buffer Saline containing 0.1% Triton-X (PBSTx) for 30 mins. Following fixation and blocking, the cells were incubated with a primary antibody cocktail containing anti-DSS1 (Catalogue# NB100-1334, Novus Biologicals) and anti-PML (Catalogue#sc-966, SCBT) for 1h at RT. After that, the cells were washed 3 times with 0.1% PBSTx for 5 mins each and incubated with a secondary antibody cocktail containing anti-goat Alexa FluorTM 647 (for DSS1), anti-mouse Alexa FluorTM 488 (for PML) and Hoechst 33342 (Catalogue# H3570, Invitrogen) for 1h at RT. Following this incubation, the cells were subjected to 3 washes, each lasting 5 mins, with 0.1% PBSTx to ensure thorough cleansing. Subsequently, z-stack imaging was performed using a Zeiss LSM800 confocal microscope with Airyscan. The above process utilized three distinct dyes to specifically label DSS1 nuclear bodies (Red), promyelocytic leukemia (PML) nuclear bodies (Green), and the nucleus (Blue). The data file was named ‘Sample Image Stack.tif’.

### 2.2 Description of VR Equipment

All screenshots of the VR environments presented in the upcoming section were captured using an HTC Vive Cosmos Elite VR System. The VR system’s Head Mounted Display (HMD) provides an approximate FoV of around 110° through two displays refreshed at a rate of 90 Hz.

## 3 METHODS

### 3.1 ColocZStats Development

High-quality 3D reconstruction or rendering is an essential prerequisite for accurately performing colocalization analysis on confocal z-stacks, as it can reveal more detail in structures, enhance users’ recognition of colocalization areas, and elevate colocalization analysis sensitivity.

Figure 1 shows two channels of the same sample dataset and their combination displayed as colored semi-transparent voxels in ChimeraX desktop, ChimeraX VR, ExMicroVR, ConfocalVR, SlicerVR, and 3D Slicer desktop separately. Although 3D Slicer supports volume rendering, before the development of ColocZStats, it could not automatically split multi-channel confocal z-stack channels and color them, nor did it provide specialized GUI widgets for separate manipulation of each channel or navigation between them. Therefore, an image processing package, FIJI (Schindelin et al., 2012), was initially utilized to save the channels as separate multi-page z-stack files. Subsequently, these files were selected and imported as scalar volumes into the ‘Volume Rendering’ module of 3D Slicer (National Alliance for Medical Image Computing, 2024) for rendering. Their colors were then manually configured to ensure consistency with the others. Currently, ChimeraX desktop and ChimeraX VR do not support applying lighting to volumetric renderings, but only to surfaces and meshes (Goddard, T.., 2024). However, adjustable highlights or shadows are necessary for showing 3D semi-translucent voxels, as this can help create refined volumetric representations and allow viewers to distinguish intricate topologies and distance relationships more easily (The American Society for Cell Biology., 2024). Unlike them, the remaining programs in Figure 1 possess the capability of adding lighting and shadows to such renderings. For example, the ‘Volume Rendering’ module of 3D Slicer provides many options for calculating shading effects, which enables finer adjustments to the rendered volume’s appearance. Regarding the rendering effects, it was observed that the quality produced by 3D Slicer desktop or SlicerVR is not inferior to that of other visualization software programs.

**Figure 1.**
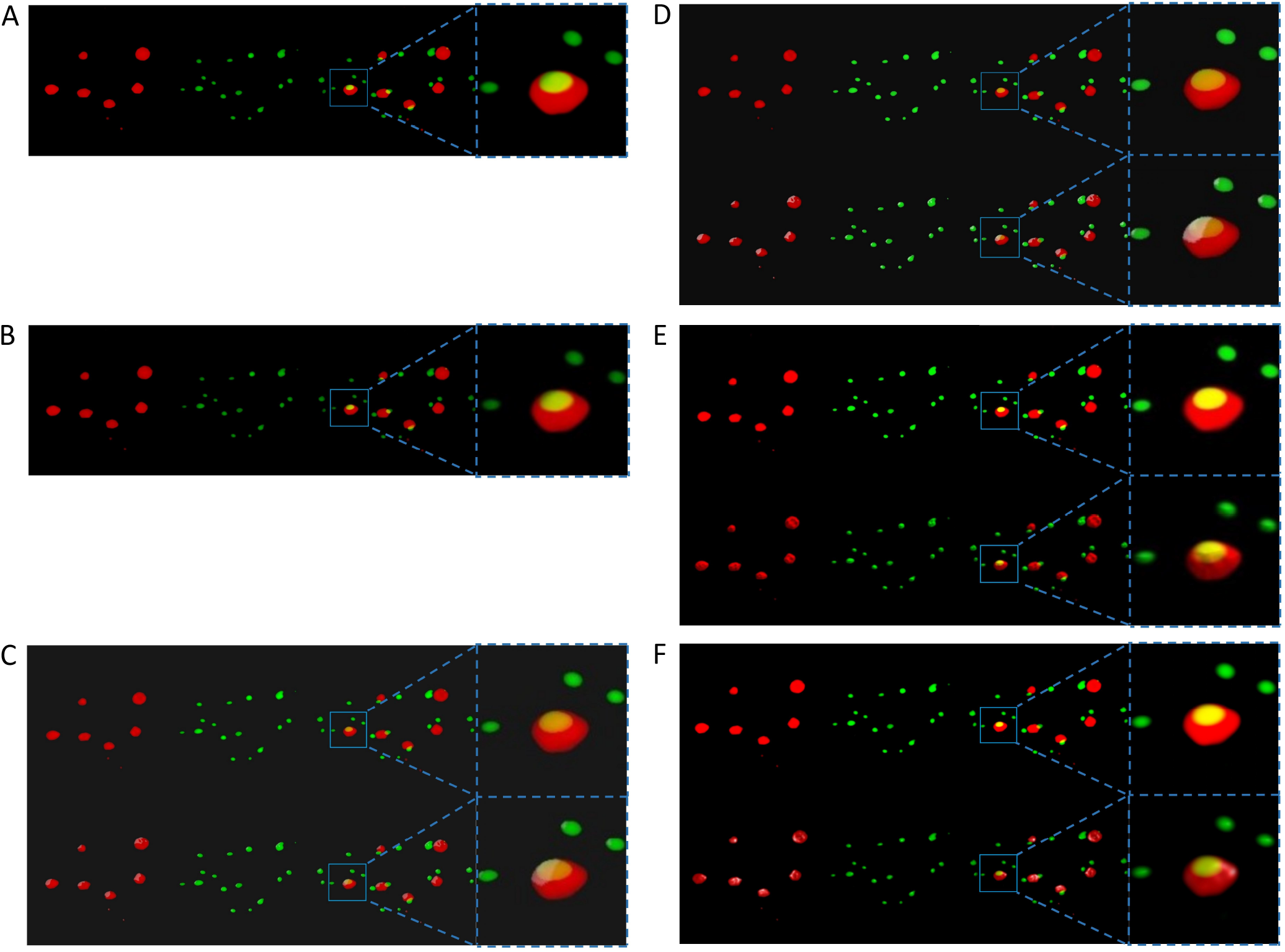
A series of screenshots displaying the visualization of the same specimen’s channels from an identical perspective across all the mentioned programs. For each sub-figure, from left to right, two separate channels of the sample stack, their superposition, and a magnification of a specific region in the superposition are shown. **(A)** The volumetric rendering of the two channels in ChimeraX’s desktop version. **(B)** The same series of scenes as **(A)**. They were captured from a VR environment created by ChimeraX VR. The first rows in **(C), (D), (E)**, and **(F)**, respectively, show the scenes without external lighting in the ExMicroVR, ConfocalVR, SlicerVR, and the desktop viewport of 3D Slicer. The second row of **(C)** to **(F)** shows the scenes with external lighting turned on.

More importantly, it has also been learned that researchers can efficiently develop, assess new methods, and add more capabilities through custom modules by benefiting from the highly extendable features of 3D Slicer. That is to say, in 3D Slicer, developers do not need extra time to redevelop primary data import/export, visualization, or interaction functionalities. Instead, they can easily call or integrate these characteristics and focus on developing new required features (Fedorov et al., 2024; Vipiana and Crocco, 2023). Although ChimeraX is also extensible, many of its functional Application Programming Interfaces (APIs) were not documented during the tool development, and many existing APIs were experimental. In contrast, 3D Slicer has provided relatively extensive API documentation and developer tutorials. Moreover, the 3D Slicer community has offered a large and active forum for developers, where considerable issues related to the tool development can be found, discussed, or raised for timely feedback. The other two, ConfocalVR and ExMicroVR, have not provided publicly available APIs. All the above factors have constituted the rationale for developing the tool for 3D Slicer.

ColocZStats is a scripted extension that utilizes the 3D Slicer APIs (Slicer Community., 2024) and is implemented using Python. The code was written in PyCharm (JetBrains s.r.o., 2024), an integrated development environment (IDE) designed explicitly for Python programming. The fundamental organizational structures of ColocZStats are ‘classes,’ which represent independent code blocks containing a group of functions and methods.

Once a multi-channel confocal z-stack is loaded into ColocZStats, in the program’s background, the stack’s metadata will be parsed to extract individual Numpy arrays of each channel. Simultaneously, several methods from a base module, ‘util.py’ and a core class, ‘vtkMRMLDisplayNode,’ of 3D Slicer will be iteratively invoked to set up separate scalar volume nodes for each channel’s Numpy array and to assign individual pseudo colors to them. Also, the rendering method from 3D Slicer’s ‘VolumeRendering’ module (Fedorov et al., 2024) will be applied to all individual channels separately, eventually presenting their merged visual appearance. Similar to most open-source medical imaging systems, such as MITK (Nolden et al., 2013), itksNAP (Yushkevich et al., 2006), and CustusX (Askeland et al., 2016), the volume rendering back-end of 3D Slicer is based on the Visualization Toolkit (VTK) (Drouin and Collins, 2018). VTK is a powerful cross-platform library that supports a variety of visualization and image-processing techniques, making it widely adopted in open-source and commercial visualization software (Bozorgi and Lindseth, 2015). The ‘Volume Rendering’ module provides three volume rendering methods: (i) VTK CPU Ray Casting, (ii) VTK GPU Ray Casting, and (iii) VTK Multi-Volume. The ‘VTK GPU Ray Casting’ is the default method for rendering because graphics hardware can significantly accelerate rendering (National Alliance for Medical Image Computing, 2024).

Moreover, at the moment a stack is loaded, a set of separate GUI elements will be created for each channel to achieve purposes such as adjusting channels’ threshold values or revealing any number of channels in the scene by controlling their visibility. The GUIs in ColocZStats are provided by the Qt toolkit (The Qt Company., 2024). Numerous modules in 3D Slicer offer a suite of reusable, modifiable GUI widgets, allowing developers to integrate them seamlessly into custom user interfaces via Qt Designer. Qt Designer is a tool for crafting and constructing GUIs with Qt Widgets. For instance, in ColocZStats, the widget integrated for controlling each channel’s threshold range is the ‘qMRMLVolumeThresholdWidget,’ which is also a widget within 3D Slicer’s ‘Volumes’ module (3D Slicer., 2024b).

By integrating and leveraging several existing classes and methods in 3D Slicer, the initial need for a series of manual operations to create appropriate volumetric representations for channels has evolved into the current state where all these steps can be automatically completed, with each channel equipped with individually controllable GUI widgets. The above process is also an illustrative example of how the high extensibility of 3D Slicer can be exploited to fulfill certain specific requirements of the tool.

### 3.2 Statistical Analysis of Colocalization

As previously mentioned in the ‘Introduction’ section, ColocZStats not only helps researchers identify the relative distribution of different cellular molecules but can also generate effective metrics for quantifying colocalization to help investigate the presence of spatial association between different channels. Next, these coefficients are described in detail.

#### 3.2.1 Pearson’s Correlation Coefficient

In the field of colocalization analysis, Intensity Correlation Coefficient-Based (ICCB) analysis methods constitute one of the primary categories of methods for assessing colocalization events. They depend on the image’s channel intensity information, which provides a powerful way to quantify the degree of spatial overlapping of two channels (Georgieva et al., 2016). A large number of colocalization analysis tools employing ICCB methods have been widely integrated into various image analysis applications (Bolte and Cordelières, 2006). Pearson’s correlation coefficient (PCC) is one of the most commonly employed ICCB methods (Bolte and Cordelières, 2006; Dunn et al., 2011). It originated in the 19th century and has been extensively used to evaluate the linear correlation between two data sets (Adler and Parmryd, 2010). When analyzing colocalization, PCC is employed to quantify the linear relationship between the signal intensities in one channel and the related values in another (Aaron et al., 2018). The PCC can be considered as a normalized assessment of two channels’ covariance (Aaron et al., 2018). The formula for PCC applied in ColocZStats is defined below, with the signal intensities of two channels at each voxel included in the calculation:

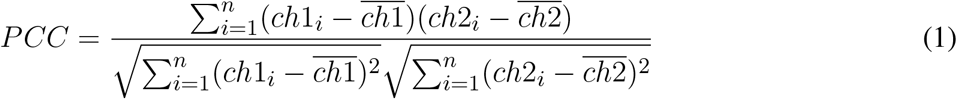

For any pair of selected channels in the tool, the *ch*1_*i*_ and *ch*2_*i*_ represent the intensity values of each channel at voxel *i*, respectively, and the 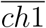 and 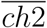 represent the mean intensities of each channel, respectively.

The range of PCC is between +1 and -1. +1 indicates that the two channels are entirely linearly correlated, and -1 means that the two channels are perfectly but inversely correlated. The value of zero means that the distributions of the two channels are uncorrelated (Dunn et al., 2011). Although PCC is theoretically not influenced by thresholds, they are incorporated into the tool’s computation to handle specified threshold settings uniformly across all coefficients computed by ColocZStats. The specific approach of setting thresholds for calculating PCC is similar to that of the ‘Colocalization Analyzer’ in the Huygens Essential software (Scientific Volume Imaging B.V.., 2024a). The Huygens Essential software (Scientific Volume Imaging B.V.., 2024b) is a widely acknowledged desktop software for the visualization and analysis of microscopic images (Pennington et al., 2022; Davis et al., 2023; Volk et al., 2022). The distinction between the two is that, in ColocZStats, not only the value of the lower threshold but also that of the upper threshold can be specified for each channel. The purpose of allowing the setting for channels’ upper threshold values is analogous to the design of ConfocalVR’s previous ‘Cut Range Max’ widget or its latest ‘Max Threshold’ widget, which is used to filter out oversaturated voxels that may be occasionally observed by observers in some particular circumstances (Stefani et al., 2018). During the calculation process, if a particular channel is set with lower and upper thresholds, voxel intensities greater than the upper threshold will be set to zero. Based on that, the lower threshold will be subtracted from the remaining voxel intensities. If any negative voxel values occur after the subtraction, they will be set to zero.

#### 3.2.2 Intersection coefficients

In distinction from the PCC, which is calculated based on actual voxel intensities, more straightforward coefficients can be calculated based only on the presence or absence of signals in a voxel, regardless of its actual intensity value. All intersection coefficients calculated by ColocZStats are examples of such coefficients, and the computation methods were borrowed from the Huygens Colocalization Analyzer (Scientific Volume Imaging B.V.., 2024a). Similarly, the difference is that the upper threshold value for each selected channel is included in ColocZStats’s calculation. To be more specific, once a voxel’s intensity value is within a specific intensity range bounded by upper and lower thresholds, it can be regarded as having some meaningful signal. If so, its value could be considered as 1, regardless of its actual intensity, and 0 otherwise. This indicates that a binary image *ch*1_*weight*_ with intensity *ch*1_*weight*,*i*_ at voxel *i* can be created based on the voxel’s actual intensity *ch*1_*i*_ and the channel’s intensity range. This explanation is illustrated by taking the first channel, *ch*1, of all selected channels as an example (the same applies to all the other channels):

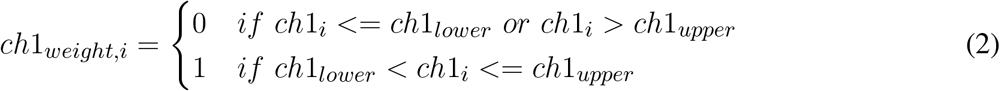

Based on the definition of *ch*1_*weight*,*i*_, another metric called the global intersection coefficient (*I*) can be calculated. It is defined as the ratio of the total volume of voxels where all channels intersect to the total volume of all channels. In other words, it calculates the proportion of voxels having valid intensity values in all thresholded channels. Multiplying this figure by 100 is interpreted as the intersection’s volume percentage. For any two specified channels in ColocZStats, the formula of *I* is defined as follows:

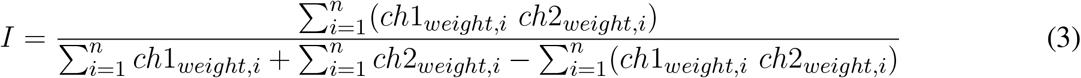

The numerator refers to the intersecting voxels’ total volume. For each voxel, the overlapping contribution is defined as the product of *ch*1_*weight*,*i*_ and *ch*2_*weight*,*i*_. The denominator represents the total volume of the two channels, which is defined as the sum of the volume of the first channel and the volume of the second channel minus their intersection’s total volume (to prevent double counting). Another two individual intersection coefficients are derived from *I*, which describes what proportion of the first and second channels are intersecting:

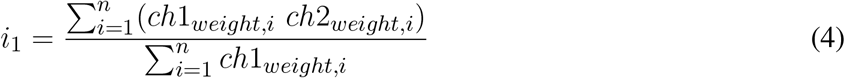

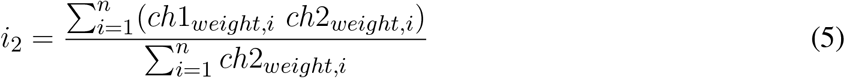

Another contribution of ColocZStats is that it allows researchers to choose up to three channels in a z-stack for statistical analysis. Extending from the above-mentioned formulas, the intersection coefficients’ formulas tailored to this scenario have also been proposed. In this case, the formula for *I* is given as:

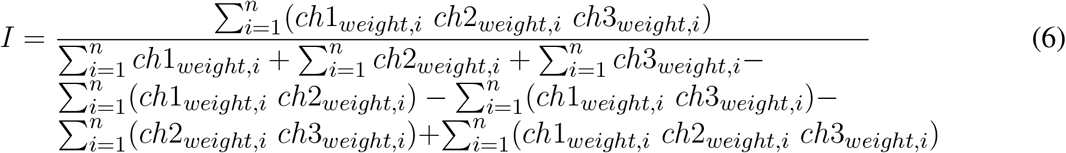

Likewise, the three channels’ intersecting volume of a given voxel *i* is defined as the product of *ch*1_*weight*,*i*_, *ch*2_*weight*,*i*_, *ch*3_*weight*,*i*_, while the sum acts as the numerator. The denominator is the total volume of the three channels, determined by the inclusion-exclusion principle for three sets (Chen, 2014). Also, three respective intersection coefficients can be obtained to exhibit the proportion of the intersection in each channel:

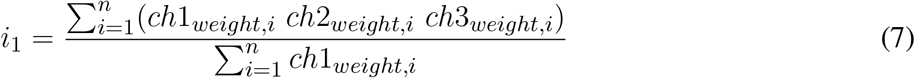

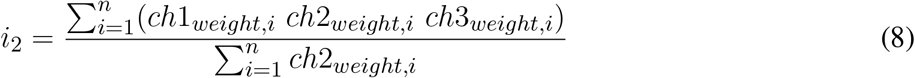

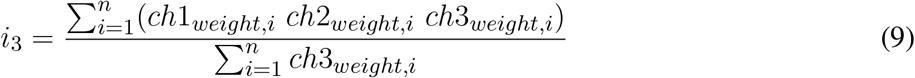

## 4 RESULTS

### 4.1 Availability of ColocZStats

ColocZStats is freely available under an MIT license and requires the stable version of 3D Slicer (3D Slicer., 2024a) for compatibility. The Slicer community maintains a website called the ‘Slicer Extensions Catalog’ for finding and downloading extensions, and ColocZStats is available there (3D Slicer., 2024c). The ‘Extensions Manager’ in 3D Slicer provides direct access to the website, facilitating easy installation, updating, or uninstallation of extensions with a few clicks in the application. As shown in Figure 2, ColocZStats can be found within the ‘Quantification’ category in the catalog. After installing the ColocZStats, it will be presented to users as a built-in extension. More information about installing and using ColocZStats can be found on the homepage of its repository on GitHub (https://github.com/ChenXiang96/SlicerColoc-Z-Stats).

**Figure 2.**
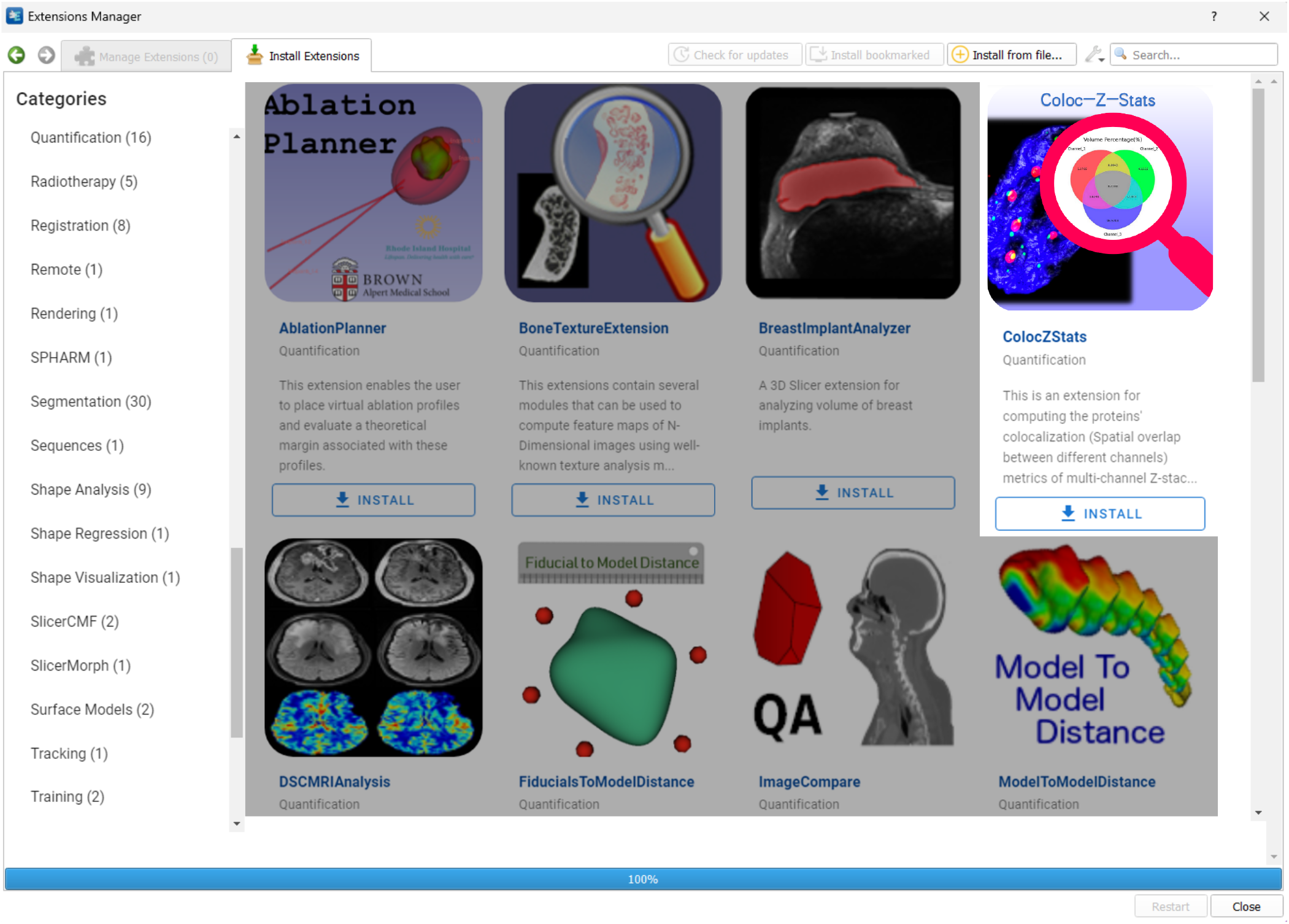
A screenshot showing the ‘Slicer Extensions Catalog’ via the ‘Extensions Manager’ in 3D Slicer. ColocZStats is offered within the ‘Quantification’ category.

### 4.2 Input Image File

For compatibility with ColocZStats, each input image file must be a 3D multi-channel confocal image z-stack in TIFF format that maintains the original intensity values, with each channel in grayscale. Each channel must possess the same dimensions, image order, and magnification. Even though ColocZStats supports loading z-stacks containing up to fifteen channels, only up to three channels can be specified for each colocalization computation because the statistical analysis and the possible interactions between channels become more complex with each additional channel.

### 4.3 User Interface

Figure 3 shows an example of ColocZStats’s user interface, which consists of two separate areas: a control panel on the left side that provides a series of interactive widgets, and a 3D view on the right side that displays the volumetric rendering of the loaded sample image stack. Users can adjust these volume-rendered images and calculate the corresponding colocalization metrics by manipulating the widgets on the control panel. To display the details of these widgets clearly, a close-up of the control panel is shown in Figure 3. The following subsection describes a typical workflow for ColocZStats.

**Figure 3.**
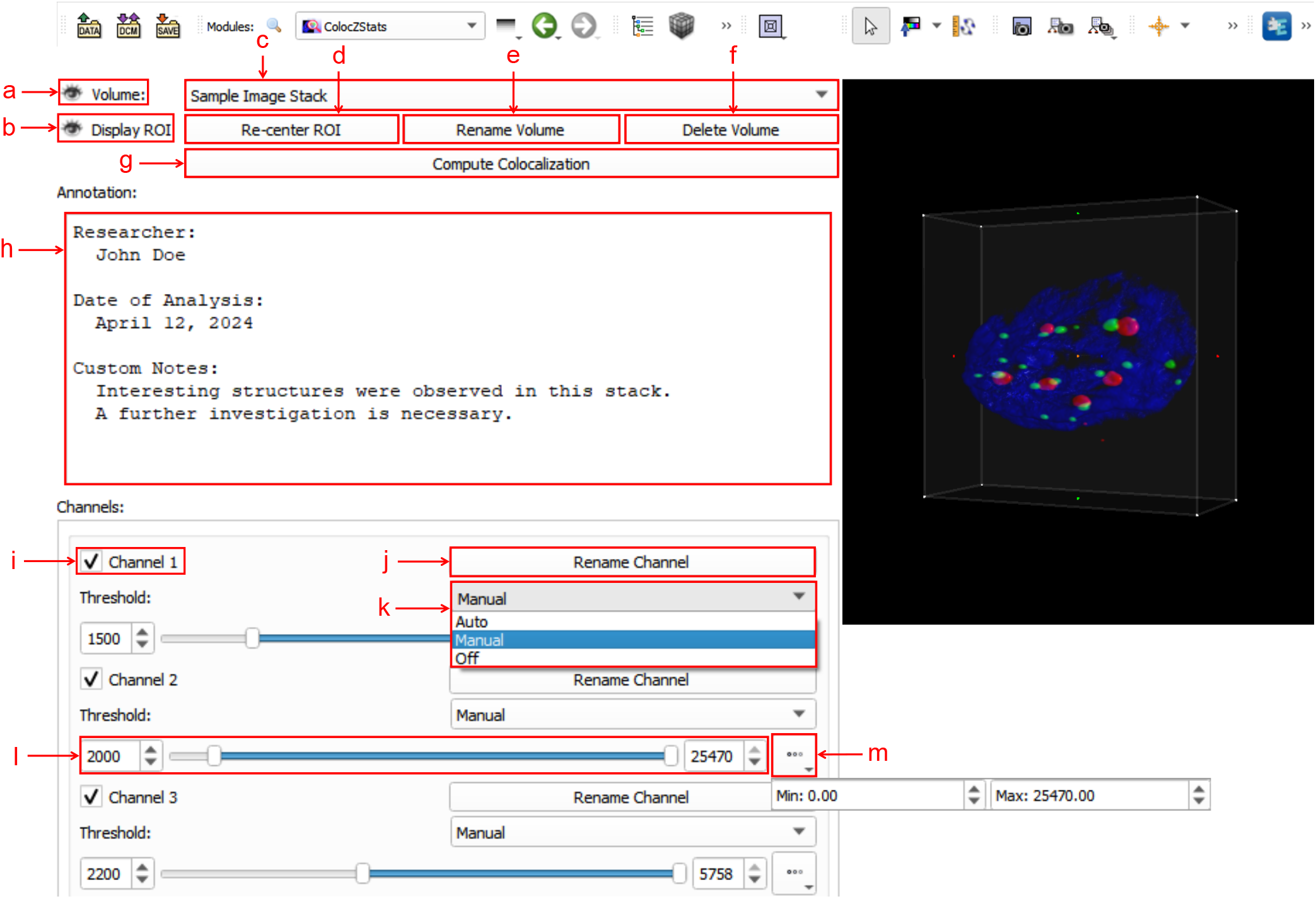
The Graphical User Interface (GUI) of ColocZStats. **(a)** A clickable eye icon for managing the visibility of the entire image stack’s rendering. **(b)** A clickable eye icon for managing the visibility of the adjustable box for selecting ROI regions. **(c)** A combo box for switching between multiple loaded image stacks, displaying the stack’s filename without extensions by default. **(d)** A button for repositioning the rendering within the ROI box to the 3D view’s center. **(e)** A button for triggering a pop-up text box for customizing the displayed name on the combo box. **(f)** A button for deleting the current stack from the scene, along with its associated annotation and GUI widgets. **(g)** A button for performing colocalization analysis. **(h)** A text field for adding a customized annotation for the current stack. **(i)** A checkbox for managing the visibility of each channel. The indices in these default channel name labels start from 1. **(j)** A button for triggering a pop-up text box for customizing the corresponding channel’s name label. **(k)** A drop-down list for selecting the channel threshold control mode. It comprises three options: ‘Auto,’ ‘Manual,’ and ‘Off’. **(l)** Adjustable sliders for setting lower and upper thresholds. Both values can also be specified in the two input fields. **(m)** A drop-down box for displaying the initial threshold boundaries of the associated channel.

### 4.4 The Workflow of ColocZStats

A typical workflow of ColocZStats is shown in Figure 4A, which is divided into five steps: (i) ‘Inputs’; (ii) ‘Visualization’; (iii) ‘Channel Selection & Thresholding’; (iv) ‘ROI Selection’; and (v) ‘Analysis.’ The sequence of certain steps in this workflow can be slightly adjusted based on particular conditions or personal preferences. In 3D Slicer, a file browser can be opened by clicking the ‘DATA’ button at the upper-left corner to load a multi-channel confocal z-stack file into the scene. All channels’ colored volume-rendered images will immediately appear in the 3D view by default when the stack is loaded into ColocZStats, and they can be moved and rotated arbitrarily with mouse clicks and movements.

**Figure 4.**
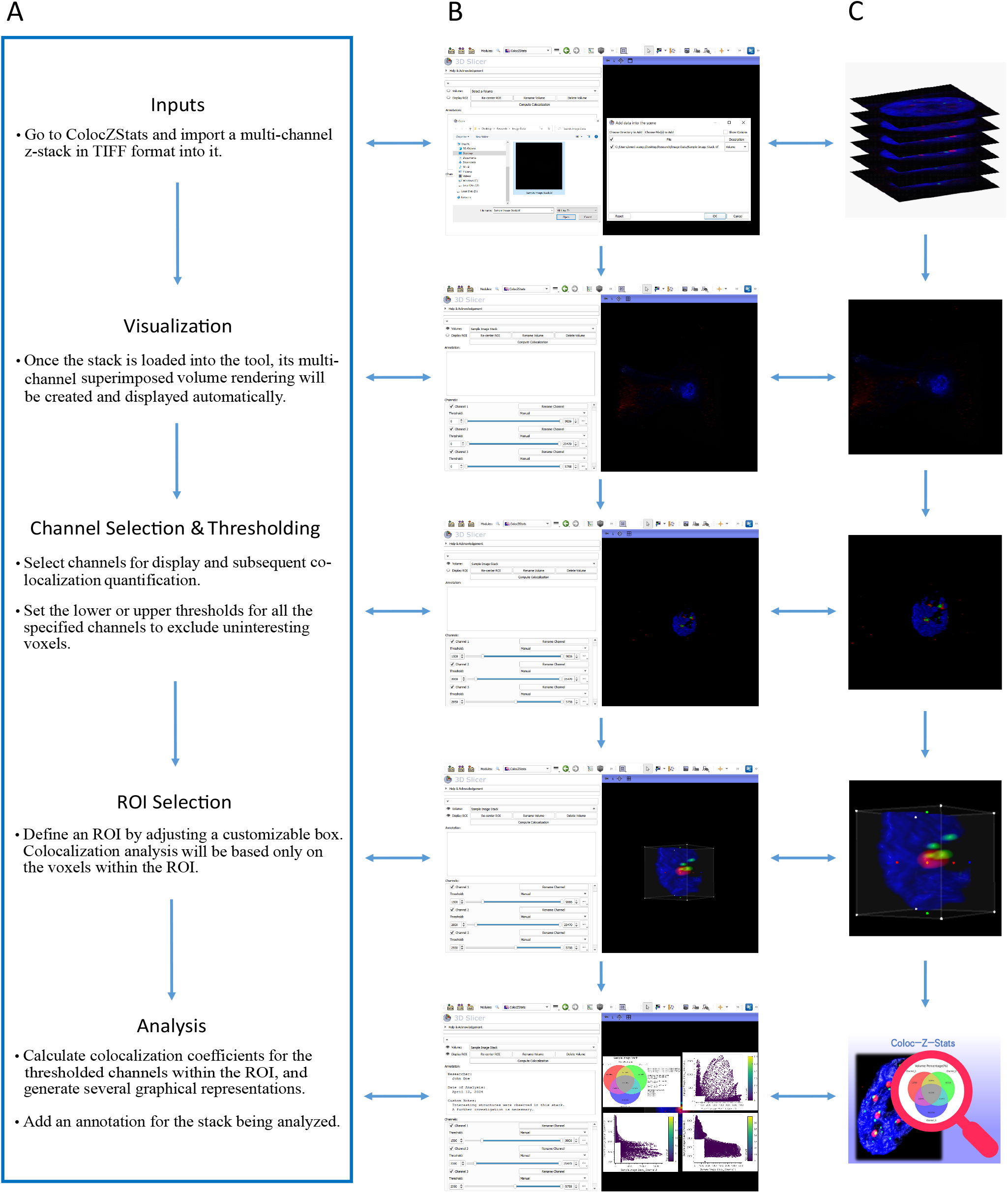
**(A)** Typical workflow of ColocZStats. **(B)** Example operation scenarios in ColocZStats that correspond to the steps in the workflow. **(C)** Visual abstractions that correspond to the steps in the workflow.

The widgets in the ‘Channels’ sub-panel will also be displayed concurrently upon loading the input file. The checkbox in front of each channel’s name label is not only used for managing the visibility of the channel’s rendered volume but also for determining whether the channel will be included in the colocalization analysis. Figure 4B,C represents an example where all three channels of the sample data are selected. After the channel selection, threshold segmentation is often necessary because it is a helpful approach for extracting interesting voxels from the image. For individual channels, as any slider for controlling the thresholds is adjusted, the display range of the channel’s volumetric representation will be changed synchronously, facilitating users’ observation and the decision-making for the following analysis. By default, when a confocal z-stack is imported, the checkboxes of all its channels are turned on, and the threshold ranges for all channels are displayed as their original ranges, respectively.

The ROI box integrated into ColocZStats facilitates researchers’ analysis of any parts of the loaded stack. By default, there is no ROI box for the imported stack in the scene. Upon the first click on the eye icon beside the ‘Display ROI’, an ROI box will be created, and after that, its visibility can be customized to be toggled on or off. While the ROI box is displayed, depth peeling will automatically be applied to the volumetric rendering to produce a better translucent appearance. The handle points on the ROI box can be dragged to crop the rendered visualization with six planes. As any handle point is dragged, the volumetric rendering outside the box will disappear synchronously. The final analysis will only include the voxels inside the ROI box. When there is a need to analyze the entire stack, the ROI box should be adjusted to enclose it completely. The functionality to extract the voxels inside the ROI is enabled by the ‘Crop Volume’ module, and the related methods will be called in the back-end when a calculation is executed. The ‘Crop Volume’ module is a built-in loadable module of 3D Slicer that allows the extraction of a rectangular sub-volume from a scalar volume. With the above control options, users can conveniently configure voxel intensity thresholds and ROI for all channels to be analyzed while intuitively focusing on the critical structures to obtain the final colocalization metrics.

The ‘Analysis’ step is performed when clicking the ‘Compute Colocalization’ button; all the thresholded channels and overlapping areas inside the ROI will be identified in the program’s background process. Subsequently, the colocalization coefficients described previously will be computed. Meanwhile, a series of graphical representations will pop up on the screen, such as a Venn diagram illustrating volume percentages and 2D histograms showing the combinations of intensities for all possible pairs of selected channels. The following subsection provides detailed descriptions of these generated graphical representations. In addition, users can add a custom annotation for the associated image stack at any step after it is loaded to record any necessary information.

### 4.5 Produced Graphical Representations

### 4.5.1 Venn Diagrams

ColocZStats allows two or three channels to be selected simultaneously to perform colocalization measurements. The Venn diagrams generated from the above two scenarios allow researchers to quickly recognize the volume percentages of overlapping regions along with that of the remaining parts. A popular Python package, ‘matplotlib-venn,’ was utilized to implement this functionality while leveraging its ‘venn2 unweighted’ and ‘venn3 unweighted’ functions for the creation of the two kinds of Venn diagrams consisting of two or three circles without area-weighting, respectively.

In the Venn diagrams generated, the colors in these circles match the colors in the volume-rendered images, and the area where all circles intersect signifies the part where all specified channels intersect, with the displayed percentage corresponding to the result obtained by multiplying the global intersection coefficient by 100 and retaining four decimal places. Based on this, the percentages of the remaining parts of these channels can be derived and displayed in the other areas of the Venn diagrams. Besides, the Venn diagram’s title matches the specified name on the GUI’s combo box, and the channel names shown correspond to the channel name labels defined on the interface.

The examples shown in Figure 5 demonstrate a typical scenario of setting lower thresholds for each channel and configuring their upper thresholds to their maximum values. Figures 5E, F are also components of the result illustrations composed together with the two Venn diagrams, respectively, and for clarity, they were extracted as two separate figures.

**Figure 5.**
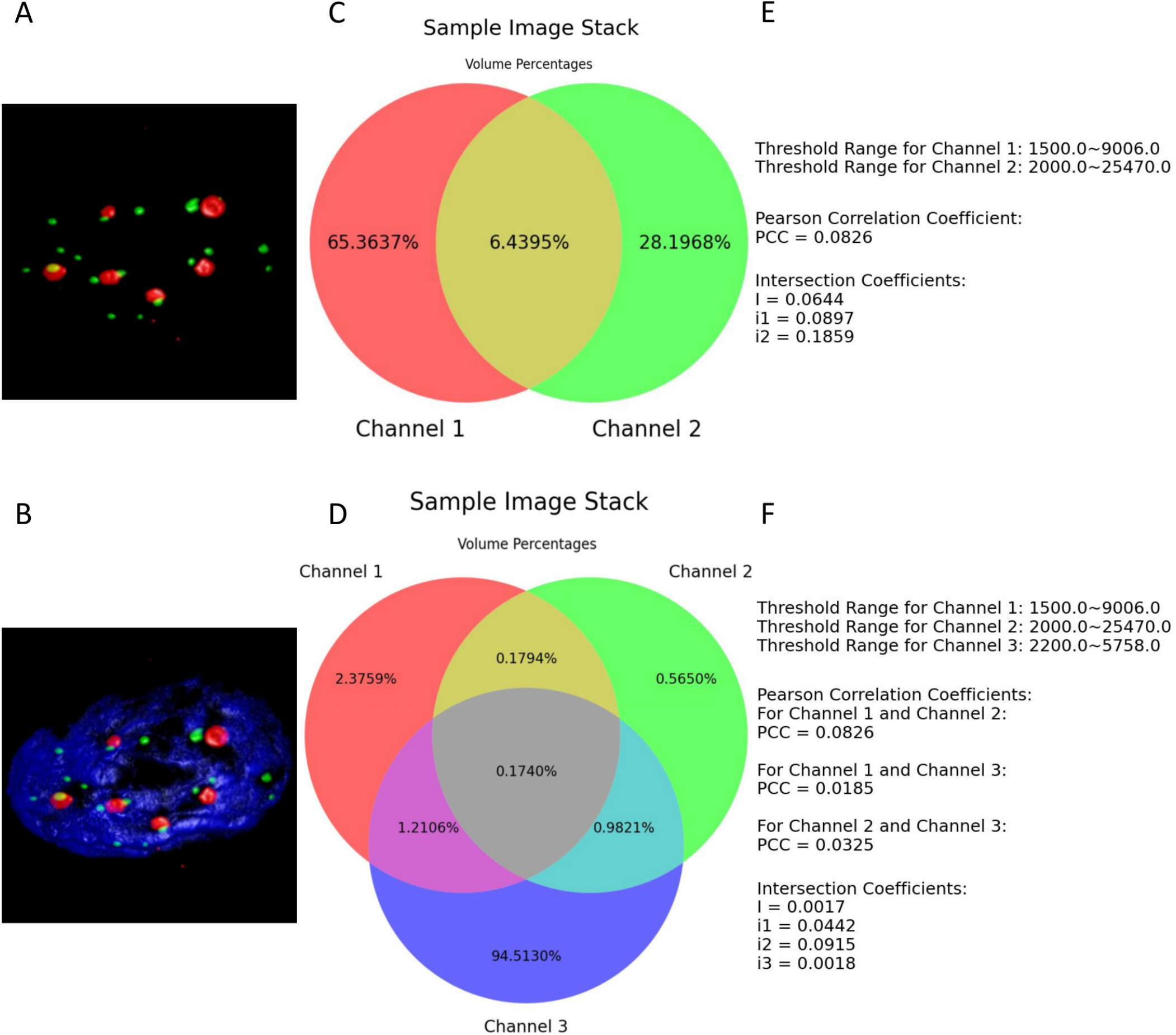
**(A)** and **(B)** present the thresholded volume renderings when two or three channels of the entire sample image stack were selected. **(C)** and **(D)** depict the respective resulting Venn diagrams. **(E)** and **(F)** illustrate all the defined thresholds and the colocalization metrics results are retained to four decimal places.

#### 4.5.2 2D Histograms

The 2D histograms produced by ColocZStas serve as a supplementary tool for visually assessing colocalization, providing a qualitative indication. The feature of generating 2D histograms has become a basic functionality of most colocalization analysis software. One specific application of 2D histograms is that they can be employed to identify populations within different compartments (Brown et al., 2000; Wang et al., 2001). For any two channels, a 2D histogram illustrates the connection of intensities between them, where the x-axis corresponds to the intensities of the first channel, and the y-axis corresponds to the intensities of the second channel. The histogram’s points can be observed closely gathering along a straight line if the two channels are highly correlated. The line’s slope indicates the two channels’ fluorescence ratio. Following the method introduced in Caltech’s ‘Introduction to Data Analysis in the Biological Sciences’ course in 2019, in ColocZStats, a Python library, called ‘Holoviews,’ (Stevens et al., 2015) was applied to plot the 2D histograms. With the scenario corresponding to Figure 5A, Figure 6 demonstrates the generation of a 2D histogram. For any precise position within the channels, the related intensities of both channels are combined to define a coordinate in the 2D histogram. Simultaneously, the count of points at this coordinate is incremented by one. As shown in Figure 6C, the blank area in the lower-left corner represents the background. For any background’s voxel, the intensity values of its two channels are both outside the respective valid channel threshold ranges, so such voxels will not be plotted as points in the 2D histogram. The definition of the valid threshold ranges mentioned in this context is consistent with those applied when calculating the intersection coefficients. The color bar on the histogram’s right side indicates the number of points with the same intensity combinations. The 2D histograms generated by ColocZStats will be saved as static images and interactive HTML files that can be viewed in more detail.

**Figure 6.**
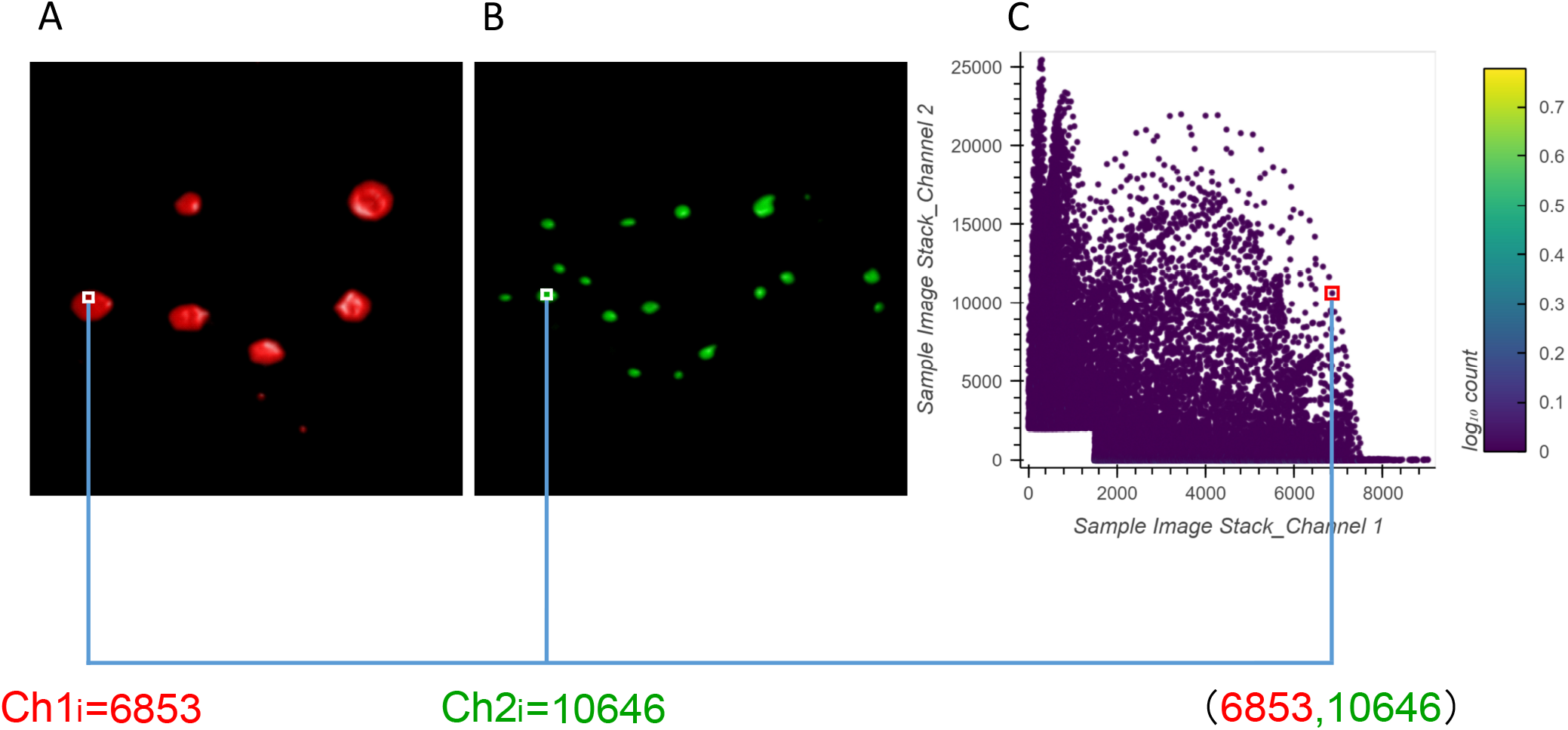
A visual explanation of a 2D histogram’s generation process. Adapted from ‘Scientific Volume Imaging-2D histograms,’ 2024, Scientific Volume Imaging B.V., https://svi.nl/TwoChannelHistogram, Copyright 1995-2024 by Scientific Volume Imaging B.V.. **(A)** The volume rendering of channel 1 and an example intensity at a specific position. **(B)** The volume rendering of channel 2 and an example intensity at the same position. **(C)** The coordinates in the 2D histogram are generated by the combination of those intensities.

The correspondence between the example 2D histogram in Figure 6 and its related Venn diagram is depicted in Figure 7. The defined threshold value boundaries for the channels delineate four regions in Figure 7A. In this 2D histogram, three colored regions, and their outlines aligned with the boundaries, are used to highlight all possible distribution positions of points. All points within the yellow region and along its yellow outlines represent all the voxels that contain valid signals in both channels, and the percentage of all these points aligns with the percentage displayed in the yellow area of Figure 7B. Likewise, the points distributed in the red or green parts represent the voxels with exclusively valid signals in channel 1 or channel 2, respectively, and the percentage of points in the two regions are consistent with the values shown in the Venn diagram’s red and green areas, respectively. Also, no points are plotted on coordinates corresponding to ‘b,’ ‘c,’ and ‘d’ or the boundaries formed between them, as they lie outside the defined threshold ranges.

**Figure 7.**
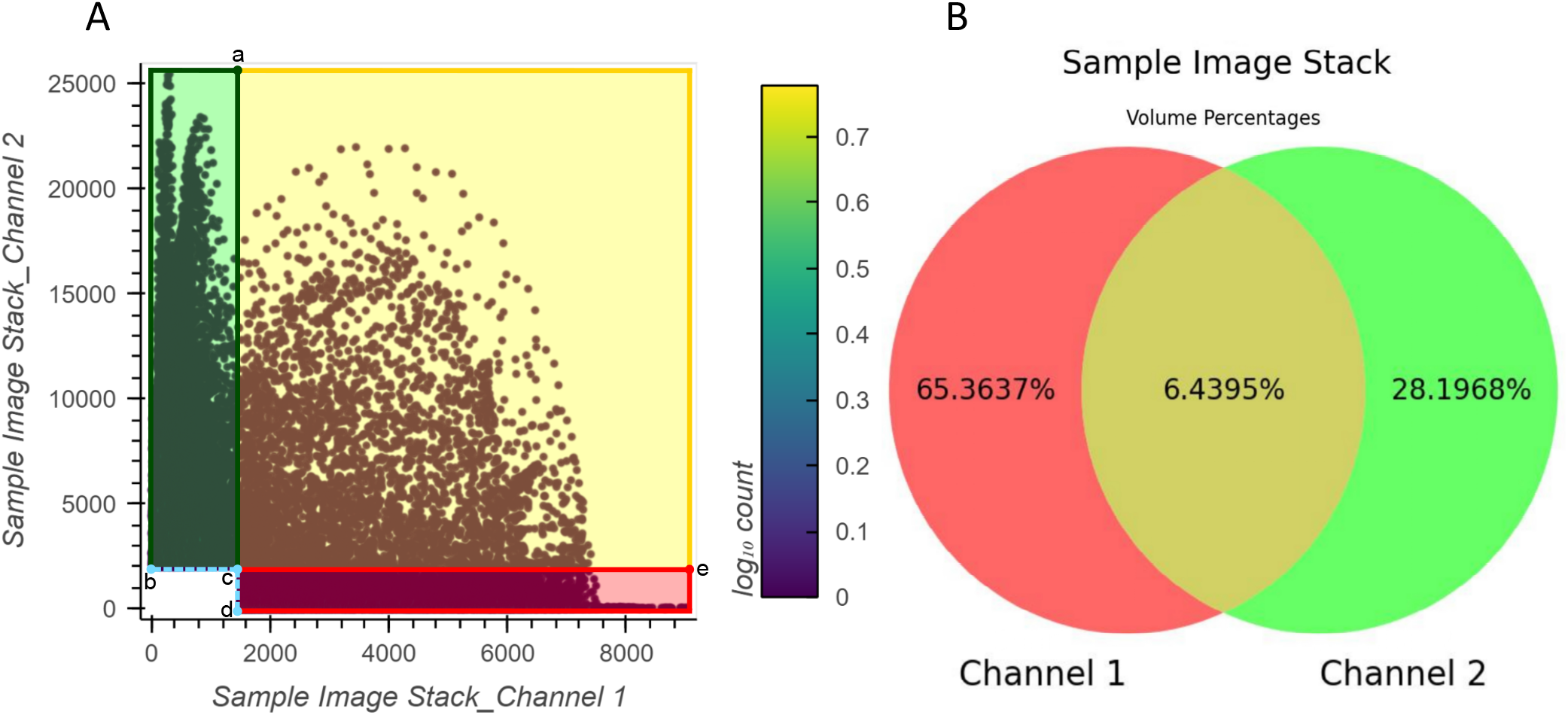
An illustration depicting the relationship between the example 2D histogram **(A)** and its corresponding Venn diagram **(B)**. Coordinate of point **(a)**: (1500,25470); Coordinate of point **(b)**: (0,2000); Coordinate of point **(c)**: (1500,2000); Coordinate of point **(d)**: (1500,0); Coordinate of point **(e)**: (9006,2000).

### 4.6 Results Spreadsheet

After clicking the ‘Compute Colocalization’ button, a comprehensive results spreadsheet will be automatically saved for researchers’ further reference or sharing. It contains all graphical representations, coefficient results, and the ROI-related information generated from each calculation. Please refer to the document/Supplementary Materials for a detailed description of the spreadsheet.

### 4.7 Case Study

This subsection provides an example demonstrating a specific scenario of applying ColocZStats to perform a colocalization analysis task. It primarily focuses on how changes in the threshold range of an individual channel influence the objective quantitative colocalization indicators and the variations in colocalization degree that can be revealed during this process. The ROI box remains unchanged throughout the four cases shown in Figure 8, and the red channel is individually assigned four distinct lower threshold values. In contrast, the threshold ranges for the other channels remain constant. By observing the 3D rendering appearance of the data provided by this tool, a continuous change can be found; that is, as the red channel’s lower threshold gradually increases, the volume of its overlap with other channels shows an evident decreasing trend. Combining a series of objective quantifications generated by ColocZStats for each case aids in validating this subjective visual impression and obtaining a more reliable assessment.

**Figure 8.**
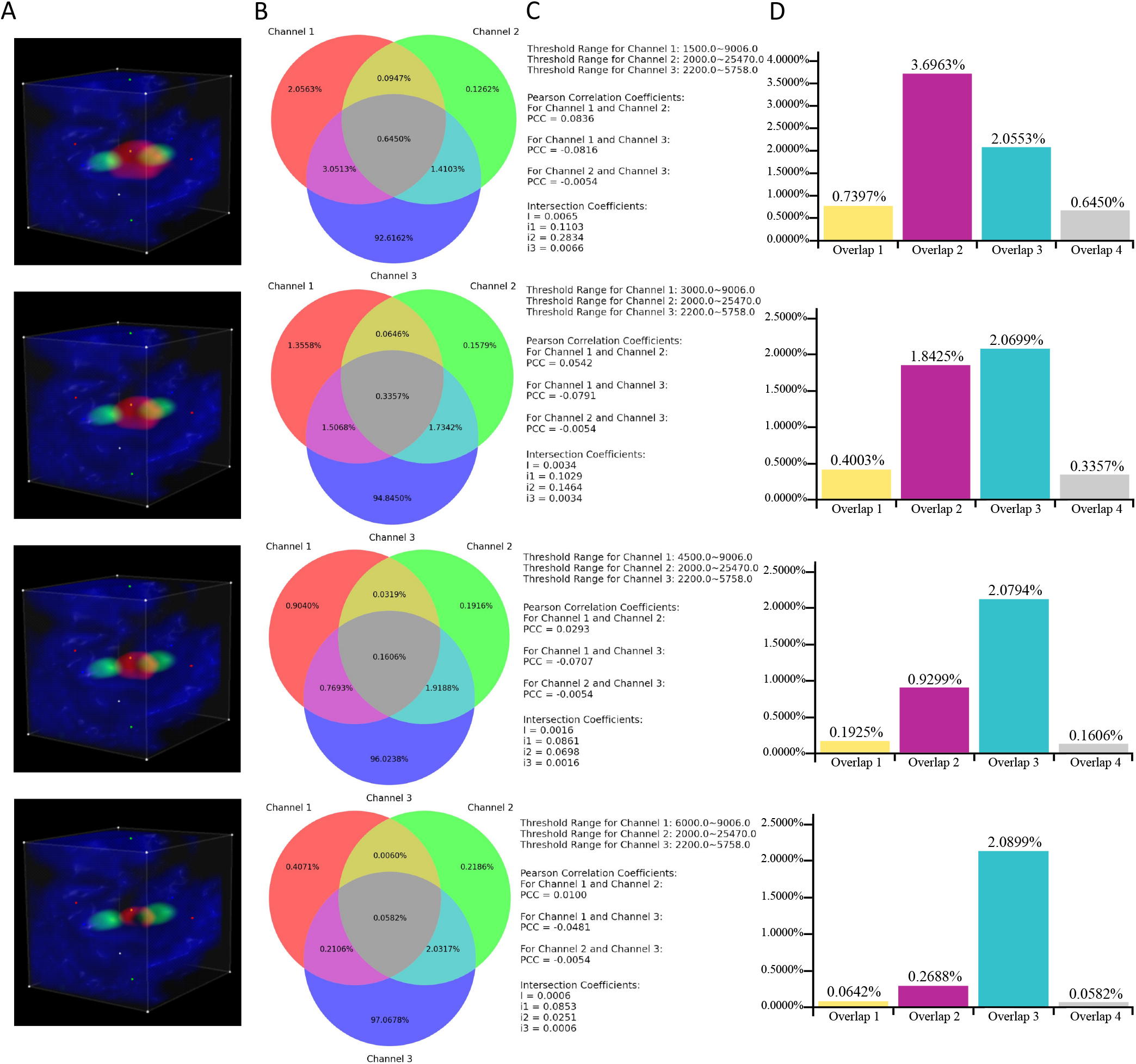
Illustration of Case Study. **(A)** Volume rendering of all channels within the same ROI box. **(B)** Venn diagrams corresponding to all the cases. **(C)** All the associated thresholds and colocalization metrics’ results. **(D)** Additional histograms illustrating the proportions of overlaps. They were created based on the data derived directly from the Venn diagrams. ‘Overlap 1’: The overlap between Channel 1 and Channel 2. ‘Overlap 2’: The overlap between Channel 1 and Channel 3. ‘Overlap 3’: The overlap between Channel 2 and Channel 3. ‘Overlap 4’: The overlap of all three channels.

Through Figure 8C, a noticeable phenomenon regarding the variation of colocalization coefficients is exposed: as the lower threshold value of channel 1 increases, the PCCs between channel 1 and the other two channels gradually approach 0, indicating a diminishing linear correlation between channel 1 and the other two channels. Meanwhile, as demonstrated by the additional histograms created based on the Venn diagrams, with the reduction in the threshold range of channel 1, the proportions of overlapping regions between channel 1 and the other two channels also decrease. Integrating the above observations effectively validates a biological fact: the reduction of channel 1 leads to a decline in its colocalization with the other two channels. In addition, the values of the global intersection coefficient and its three derived coefficients also gradually decrease, implying a decline in the overall colocalization degree of the three channels. This example also exhibits how the visual representations and coefficients produced by this tool can provide multiple perspectives for assisting colocalization analysis.

### 4.8 Comprehensive Comparison of Visualization Tools

The related features of the programs mentioned in the ‘Introduction’ section, including ColocZStats, for visualizing confocal z-stacks and measuring their colocalization are summarized in Table 1. This table divides these features into four categories, encompassing the most meaningful comparable features for the execution of colocalization analysis. Consequently, although certain programs may have many other functionalities, they are not included in this table. Regarding the choice of programs, ChimeraX VR and SlicerVR are VR extensions of ChimeraX and 3D Slicer correspondingly, and their image processing and analysis capabilities almost entirely depend on the specifics of each platform. As a result, only ChimeraX and 3D Slicer are listed in this table for comparison. For each feature in the table, if the programs themselves or any plugins or modules they incorporate offer matching functionality, the corresponding cell is marked with a check. For instance, in the case of 3D Slicer, its ‘Volume Rendering’ module supports loading multi-page Z-stack files; thus, the corresponding cell is checked. Another module in 3D Slicer, ‘ImageStacks,’ allows loading the image sequence of z-stack slices, resulting in the respective cell being checked as well. It is worth noting that because the ‘ImageStacks’ module is designed specifically for working with image stacks, all the listed image formats in the ‘Supported image format’ for 3D Slicer are consistent with those supported by the ‘ImageStacks’ module.

**Table 1.**
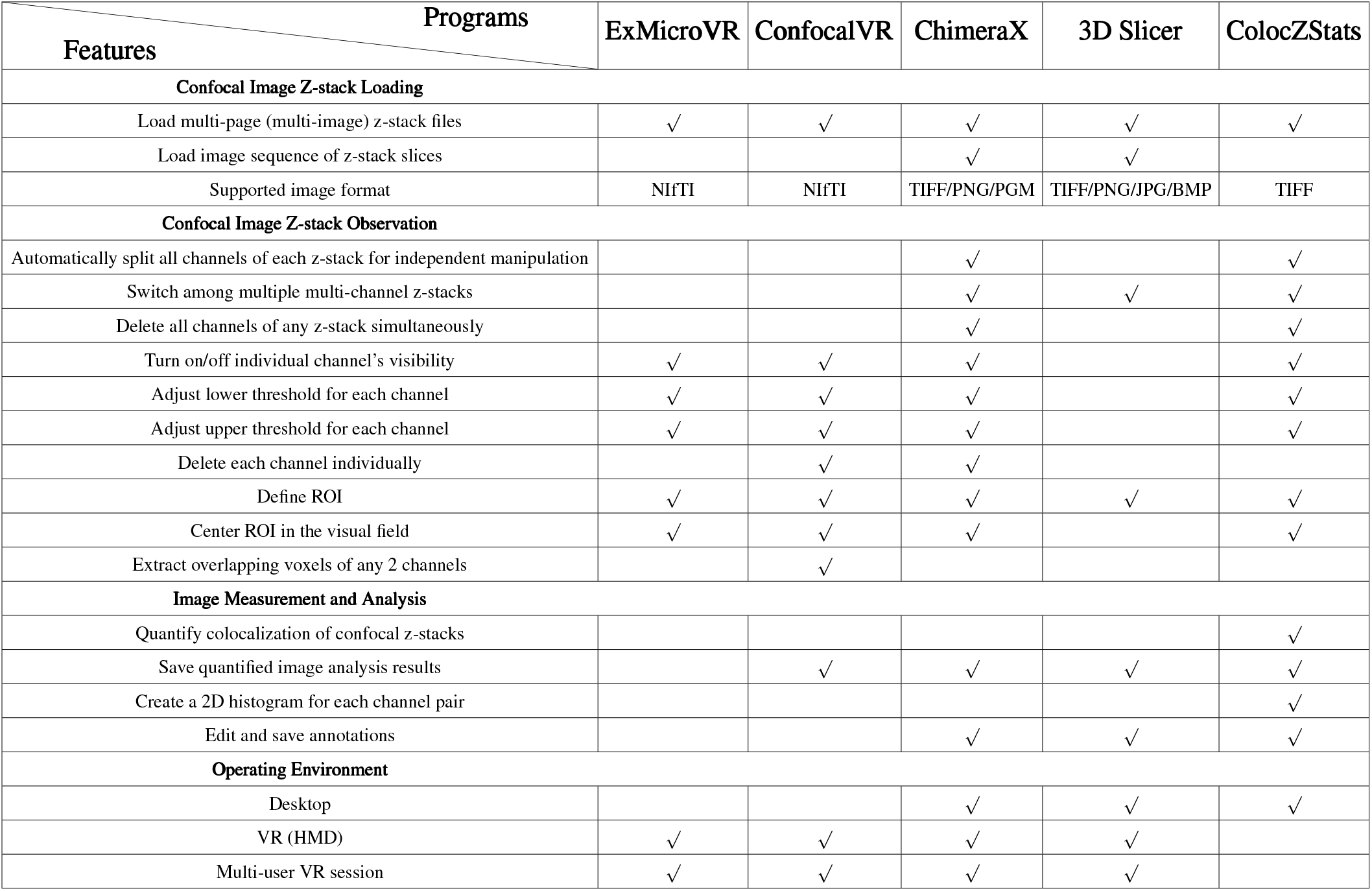
Programs Features Overview.

The category ‘Confocal Image Z-stack Observation’ summarizes the general practical features for meticulously observing the channels in confocal z-stacks. Most of them are common to several tools in this comparison. Among these, as for ‘ExMicroVR’ or ‘ConfocalVR,’ if one intends to manipulate individual channels separately, a necessary step is to extract each channel of the original z-stack as a single multi-page z-stack file using FIJI and save all of them into a designated directory for loading. Therefore, while both tools support users in manipulating channels individually, they do not have the functionality to automatically separate all channels of the z-stack to enable this operation. Before the completion of ColocZStats development, as indicated by the information from the 3D Slicer community, 3D Slicer still needed specialized tools for properly visualizing multi-channel confocal z-stacks and manipulating channels separately. Therefore, in the table, no features related to operating z-stack channels corresponding to 3D Slicer have been checked. On a side note, in the context of the third and seventh items within this category, ‘Delete’ refers to removing channels’ rendering and matching GUI widgets from the scene rather than deleting any associated files from the file system.

From the information presented in Table 1, it is indicated that in contrast to other platforms, ColocZStats not only retains several necessary control options for channels of confocal z-stacks, but can also objectively analyze the colocalization in such stacks. Significantly, this comparison also reveals some limitations in the current functionalities of ColocZStats. The current prominent limitations of ColocZStats and potential directions for future enhancements are discussed in detail in the ‘Discussion’ section.

## 5 DISCUSSION

Through a comprehensive examination of several widely recognized biological image visualization applications, all with VR functionalities and some of which already own extensive analytical features, it has been identified that these tools lack dedicated built-in options and user interfaces for 3D graphics to perform the quantitative colocalization analysis for multi-channel z-stacks generated by confocal microscopes. The development of an open-source 3D Slicer extension named ColocZStats has contributed to expanding potential solutions for addressing this challenge. Certain functionalities from multiple modules within 3D Slicer were reasonably utilized and integrated by ColocZStats, enabling the effective presentation of confocal microscopy images’ merged multi-channel volumetric appearances. This endeavor aims to support researchers in quickly discerning spatial relationships between molecular structures within organisms. On top of that, ColocZStats can generate colocalization metrics for thresholded channels within ROIs of samples, which has positive significance for biologists who now avail of a tool to gain detailed and objective insights into biological processes.

More specifically, ColocZStats allows users to select up to three channels of each Z-stack concurrently for analysis. It permits customized control over channels’ lower or upper threshold limits according to researchers’ specific needs and obtaining metrics such as PCCs for all possible channel pairwise combinations and intersection coefficients. The above distinctive characteristics distinguish ColocZStats from most tools, which typically restrict users from performing colocalization analysis between two channels at a time and only allow setting the lower threshold for each channel. Via the Venn diagram it generates, the ratios of all parts, including the part where all channels intersect, can be clearly displayed. Moreover, users can further enhance their understanding of the intensity relationship between different channels through the generated 2D histograms. For each calculation, all the results and diagrams are conveyed to researchers through a supplementary spreadsheet, making it easy for them to share or compare data. In summary, with the ultimate goal of merging with SlicerVR, ColocZStats is presently functioning as a desktop extension for 3D Slicer, incorporating an intuitive GUI that allows users to customize ROIs and define the threshold ranges for all stack channels while supporting the one-click generation and saving of colocalization analysis results. More importantly, the development of the ColocZStats extension has further enhanced the comprehensiveness of 3D Slicer, which means that the extensive audience of 3D Slicer can now seamlessly perform colocalization analysis for confocal stacks without frequently switching between different tools.

At present, as a purpose-built tool for colocalization analysis, ColocZStats requires prioritized improvements in the following aspects. Firstly, there is a need to expand the variety of colocalization metrics. In addition to the existing PCC, other coefficients belonging to the ICCB methodology that are also extensively employed, such as Manders’ coefficients, Spearman’s coefficient, and the overlap coefficient, could be integrated into the extension to further enhance its analytical capabilities. As more colocalization coefficients become incorporated into ColocZStats in the future, enhancing the capability of image pre-processing becomes increasingly essential. The need for this enhancement arises from the potential impact of excessive noise in microscopic images, which affects the correlation between distinct signals and leads to an underestimation of colocalization analysis results (Adler et al., 2008). Deconvolution is a well-established method for image filtering and restoration (Landmann and Marbet, 2004). Integrating this technique into the tool could significantly help eliminate image noise, improve image quality (Wu et al., 2012), and consequently improve the accuracy of subsequent analysis.

Meanwhile, to further increase the coverage of ColocZStats for a wider range of input images and the efficiency of analyzing multi-channel image stacks, it is anticipated that features allowing compatibility with stacks containing more channels and permitting the simultaneous selection of more channels in a single computation will be implemented.

Moreover, as elucidated in the aforementioned ‘User Interface’ subsection, ColocZStats supports changing channels’ thresholds by manually adjusting the sliders or entering values in the input fields to help biologists customize structures of interest. Nonetheless, relying on visual inspection of images to estimate suitable thresholds can be challenging and may result in inconsistent outcomes. Figure 3(k), as one of the components of the ‘qMRMLVolumeThresholdWidget’ provided by 3D Slicer, although it includes an ‘Auto’ option that can be used to assign a threshold range for any channel automatically, incorporating more reliable, robust, and objective automated threshold methods will undoubtedly further enhance the functionality of ColocZStats. The method developed by Costes et al (Costes et al., 2004). to determine the appropriate threshold value for background identification is expected to be appropriately integrated into ColocZStats. The Costes method is founded on the linear fitting of channels’ 2D histograms (Costes et al., 2004; Pike et al., 2017) and has been proven to be a robust and reproducible approach that can be readily automated (Dunn et al., 2011).

Enabling users to utilize ColocZStats within an immersive VR environment is a primary objective for future work. At the time of writing, integrating arbitrary interactive Qt widgets into the VR environment is an ongoing development effort by the SlicerVR development team. Once this feature is fully implemented, this would imply that any functionality in 3D Slicer and its extensions could be easily accessed in VR through these virtual widgets (NA-MIC Project Weeks., 2024; Pinter et al., 2020). This work illuminates the significant potential for extending ColocZStats into the immersive scene offered by Slicer VR, which will provide valuable assets and support for this forthcoming endeavor. Through these and other improvements along these lines, ColocZStats will be continually enhanced to become a more efficient and flexible software tool.

## Supporting information

Supplementary Info Sheet

Sample Spreadsheet

## AUTHOR CONTRIBUTIONS

T.B. and O.M. conceived and designed the study, obtained funding, supervised the study, and revised the manuscript; X.C. implemented the program, distributed the program to 3D Slicer, analyzed the data, prepared figures, and wrote the manuscript; T.T and A.D.J provided the confocal microscopy data. All authors have read and approved the submitted version of the manuscript.

## ACKNOWLEDGMENTS

We thank the Memorial University of Newfoundland’s School of Graduate Studies, the Memorial University of Newfoundland’s Department of Computer Science, and the Memorial University of Newfoundland’s Faculty of Medicine for their financial and equipment support.

## DATA AVAILABILITY STATEMENT

The confocal microscopy data presented in this study are included in the document/Supplementary Materials.

