## Supplementary Info Sheet for "ColocZStats: A Z-Stack Signal Colocalization Extension Tool for 3D Slicer"

### Supplementary Material

#### 1 DESCRIPTION OF SUPPLEMENTARY DATA

All loaded data in 3D Slicer can be stored in a data repository called the 'MRML scene'. All those graphical representations and statistical coefficients generated by ColocZStats, as shown in sections 4.5 and 4.7 of the article, can be obtained by loading corresponding files with the '.mrml' file extension into the current stable version of 3D Slicer (V5.6.2). In the supplementary materials, there are two folders containing MRML files, each of which is archived within a separate zip file.

Specifically, the MRML file stored in 'Scenario 1.zip' can be utilized to generate all result illustrations and statistical coefficients corresponding to sections 4.5 in the article. The MRML file saved in 'Scenario 2.zip' can be used to produce the relevant illustrations presented in section 4.7 of the article.

Additionally, an example results spreadsheet (Sample Image Stack Statistics.xlsx) is included in the supplementary materials, and Figure S1 showcases all of its sub-sheets. Among them, Figure S1D displays some crucial information related to the defined ROI. The information is automatically and selectively extracted from a JSON file containing all details of the ROI node, and the JSON file is for reloading the ROI box. By invoking the 3D Slicer's built-in exporting method for such JSON files in ColocZStats' background process, the JSON file will also be automatically saved after each calculation. Notably, the JSON file can also be saved as one of the batch files produced by clicking the 'SAVE' button of 3D Slicer. All the files included in 'Scenario 1.zip' or 'Scenario 2.zip' are the batch files generated this way, and the MRML files refer to all the other batch files in their respective folder for reloading operation scenarios.

Moreover, the images used in this study show co-localisation of DSS1 nuclear bodies (Red, 647 channel) with PML nuclear bodies (Green, 488 channel) in RMG-I cell line. Cellular nucleus (Blue, 405 channel) was stained using Hoechst 33342 (see Table S1).

Z-stack image and scenario files can be obtained from the Zenodo repository (10.5281/zenodo.11372183).

#### 2 STEPS TO LOAD MRML FILES

Before loading any MRML files into 3D Slicer 5.6.2, a necessary step is installing ColocZStats. For detailed instructions on how to install ColocZStats, please see its repository homepage on GitHub.

| Target | Primary Antibody | Secondary Antibody | Confocal Channel |
|---|---|---|---|
| **DSS1** | Anti-DSS1 Cat# NB100-1334, Novus Biologicals | Anti-goat Alexa Fluor$^{TM}$ 647 Cat# A-21447, Invitrogen | 647 |
| **PML** | Anti-PML Cat#sc-966, SCBT | Anti-mouse Alexa Fluor$^{TM}$ 488 Cat# A-11059, Invitrogen | 488 |
| **Nucleus** | Hoechst 33342 Cat# H3570, Invitrogen | | 405 |

**Table S1.** Immunocytochemistry

Taking ‘Scenario 1.zip’ as an example, please follow the steps below to load the MRML files:

1. Unzip ‘Scenario 1.zip’ to get a folder with the same name.
2. Please follow the numerical order as indicated in Figure S2 to load the ‘Scenario 1.mrml’ file from the folder obtained in the first step into the scene.

Upon loading any unzipped MRML files into 3D Slicer, all of its corresponding GUI elements and volume renderings will appear. Following this, please use the control panel of ColocZStats to select the channels and threshold ranges to be analyzed. Then, click the ‘Compute Colocalization’ button to generate the corresponding Venn diagrams, 2D histograms, and coefficient results described in the article’s respective sections. Simultaneously, all the generated diagrams and results spreadsheet will be automatically saved to the ‘Default scene location’ of 3D Slicer. Figure S3 illustrates how to specify the ‘Default scene location’ in 3D Slicer. By default, the ‘Default scene location’ is the installation location of 3D Slicer.

A

| Channels | Threshold Ranges |
|---|---|
| Sample Image Stack_Channel 1 | 1500.0~9006.0 |
| Sample Image Stack_Channel 2 | 2000.0~25470.0 |
| Sample Image Stack_Channel 3 | 2200.0~5758.0 |

B

| Timestamp |
|---|
| Fri Apr 12 21:55:02 2024 |

C

| Annotation |
|---|
| Researcher:<br>John Doe<br><br>Date of Analysis:<br>April 12, 2024<br><br>Custom Notes:<br>Interesting structures were observed in this stack.<br>A further investigation is necessary. |

H

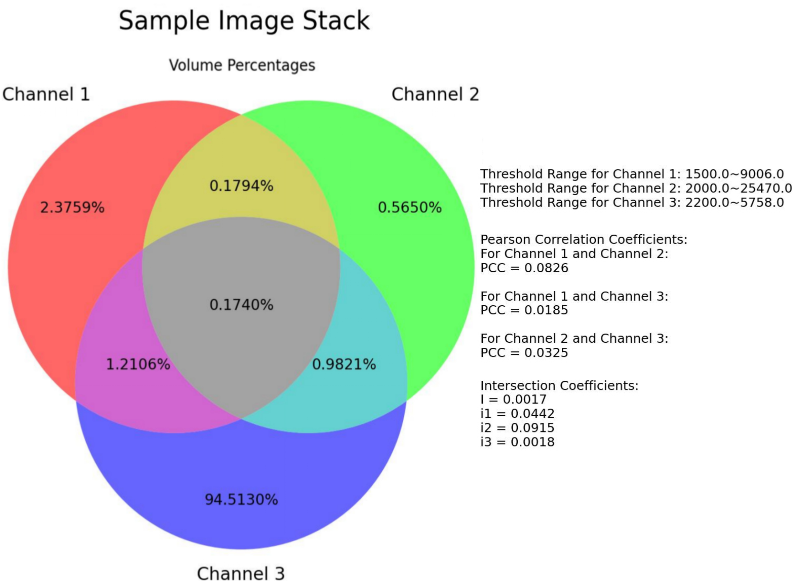


D

| | |
|---|---|
| Coordinate System: | LPS |
| Center: | [-879.5, -879.5, 50.0] |
| Orientation: | [-1.0, -0.0, -0.0, -0.0, -1.0, -0.0, 0.0, 0.0, 1.0] |
| Size: | [1760.0, 1760.0, 105.0] |
| ROI JSON File Location: | C:/Users/smed-xiangc/AppData/Local/slicer.org/Slicer 5.6.2/Sample Image Stack ROI Information.mrk.json |

E

| Channel Pairs | Pearson Correlation Coefficients (PCCs) |
|---|---|
| Sample Image Stack_Channel 1 and Sample Image Stack_Channel 2 | 0.0826 |
| Sample Image Stack_Channel 1 and Sample Image Stack_Channel 3 | 0.0185 |
| Sample Image Stack_Channel 2 and Sample Image Stack_Channel 3 | 0.0325 |

F

| Global Intersection Coefficient (I) | i1 | i2 | i3 |
|---|---|---|---|
| 0.0017 | 0.0442 | 0.0915 | 0.0018 |

G

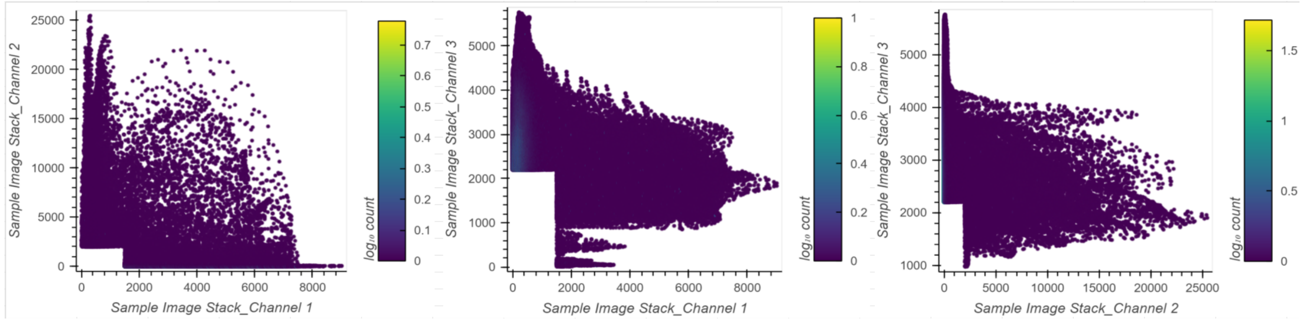


**Figure S1.** All of the sub-sheets in each spreadsheet result from each single computation. **(A)** The selected channels and their respective threshold ranges. **(B)** The timestamp of this computation. **(C)** The custom annotation for the specified image stack. **(D)** The related information of the defined ROI for this computation. **(E)** The PCCs for all possible pairwise combinations of the selected channels. **(F)** All the resulting intersection coefficients. **(G)** All the resulting 2D histograms, with each histogram illustrating a pair of channels. **(H)** The resulting illustration embedded with the Venn diagram and all coefficient results.

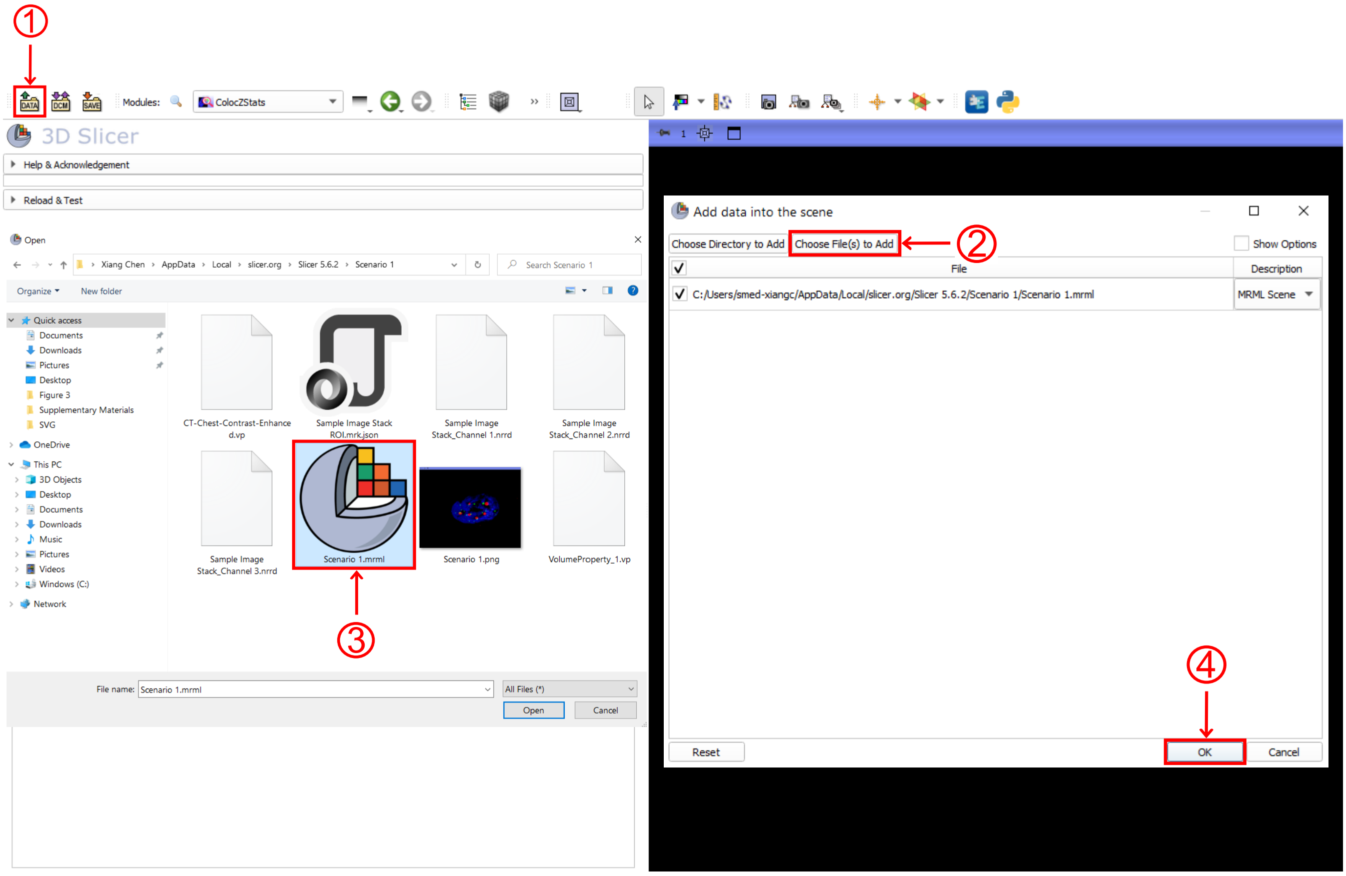


**Figure S2.** An illustration of loading ‘Scenario 1.mrml’ into ColocZStats. **(1)** Navigate to the ColocZStats extension and click on the ‘DATA’ button at the upper-left corner to open a pop-up window called ‘Add data into the scene.’ **(2)** Click the ‘Choose File(s) to Add’ button to open a file browser. **(3)** Navigate to the unzipped folder of ‘Scenario 1.zip’and select the ‘Scenario 1.mrml.’ **(4)** Click the ‘OK’ button to load the MRML file into the scene.

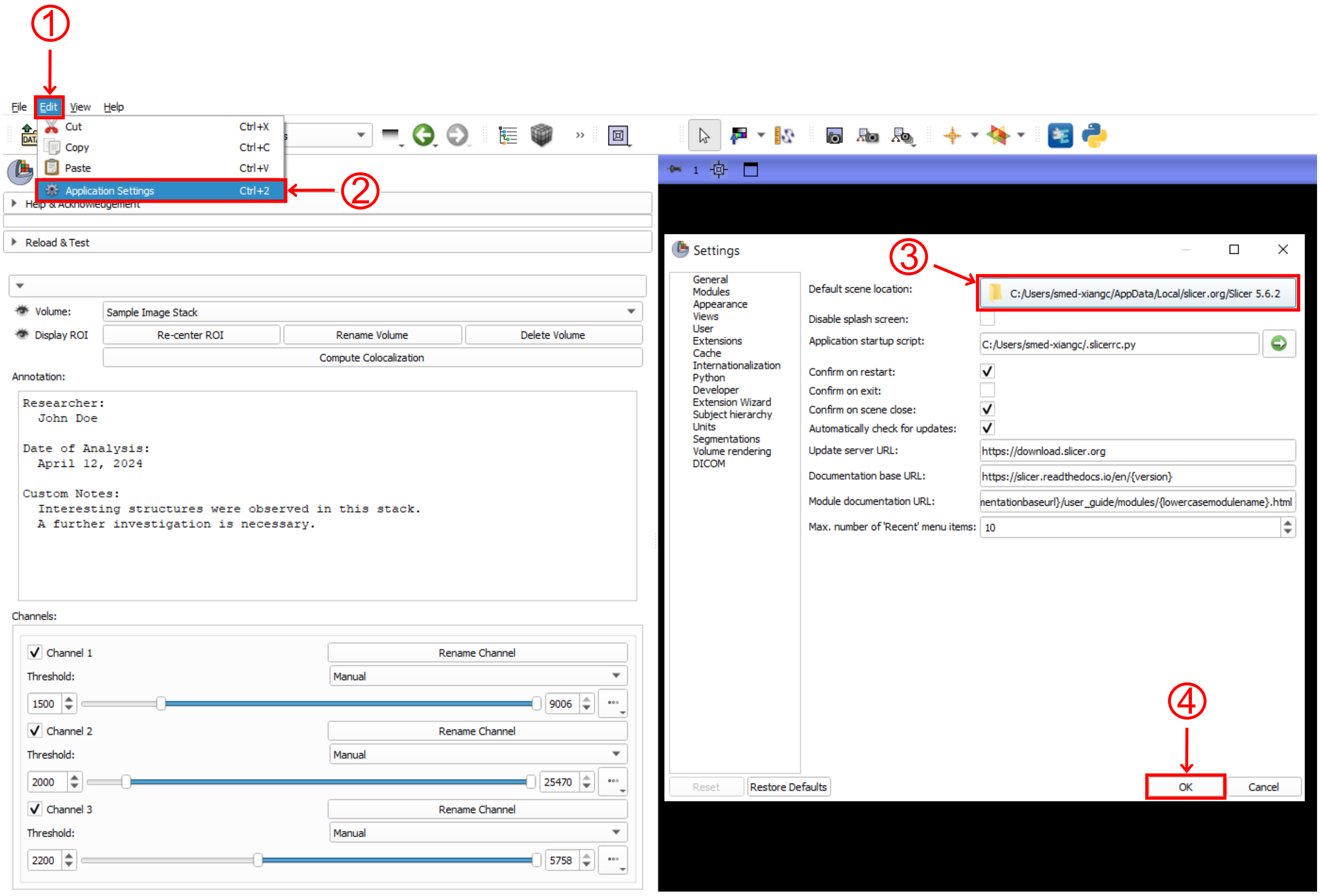


**Figure S3.** An illustration of specifying 'Default scene location' in 3D Slicer. **(1)** Click the 'Edit' button at the upper-left corner to open a drop-down list. **(2)** Click on the 'Application Settings' to open its pop-up window. **(3)** Click the button corresponding to the'Default scene location: ' to select a location with read and write permissions. **(4)** Click the 'OK' button to finish this setup.
